# Ultra-high field imaging of hippocampal neuroanatomy in first episode psychosis demonstrates receptor-specific morphometric patterning

**DOI:** 10.1101/829630

**Authors:** Min Tae M. Park, Peter Jeon, Ali R. Khan, Kara Dempster, M. Mallar Chakravarty, Jason P. Lerch, Michael Mackinley, Jean Théberge, Lena Palaniyappan

**Affiliations:** Department of Psychiatry, Schulich School of Medicine and Dentistry, Western University, London, Canada; Department of Medical Biophysics, Western University, London, Canada; Robarts Research Institute, Western University, London, Canada; Lawson Health Research Institute, London, Canada; Departments of Psychiatry and Biological and Biomedical Engineering, McGill University, Montreal, Canada; Cerebral Imaging Centre, Douglas Mental Health University Institute, Montreal, Canada; Program in Neurosciences and Mental Health, The Hospital for Sick Children, Toronto, ON, Canada; Department of Medical Biophysics, University of Toronto, Toronto, ON, Canada; Wellcome Centre for Integrative Neuroimaging, University of Oxford, Oxford, UK

## Abstract

**Objective:** The hippocampus is considered a putative marker in schizophrenia with early volume deficits of select subfields. Certain subregions are thought to be more vulnerable due to a glutamate-driven mechanism of excitotoxicity, hypermetabolism, and then degeneration. Here, we explored whether hippocampal anomalies in first-episode psychosis (FEP) correlate with glutamate receptor density via a serotonin receptor proxy by leveraging structural neuroimaging, spectroscopy (MRS), and gene expression.

**Methods:** High field 7T brain MR images were collected from 27 control, 41 FEP participants, along with 1H-MRS measures of glutamate. Automated methods were used to delineate the hippocampus and atlases of the serotonin receptor system were used to map receptor density across the hippocampus and subfields. We used gene expression data from the Allen Human Brain Atlas to test for correlations between serotonin and glutamate receptor genes.

**Results:** We found reduced hippocampal volumes in FEP, replicating previous findings. Amongst the subfields, CA4-dentate gyrus showed greatest reductions. Gene expression analysis indicated 5-HTR1A and 5-HTR4 receptor subtypes as predictors of AMPA and NMDA receptor expression, respectively. Volumetric differences in the subfields correlated most strongly with 5-HT1A (R=0.64, p=4.09E-03) and 5-HT4 (R=0.54, p=0.02) densities as expected, and replicated using previously published data from two FEP studies. Measures of individual structure-receptor alignment were derived through normative modeling of hippocampal shape and correlations to receptor distributions, termed Receptor-Specific Morphometric Signatures (RSMS). Right-sided 5-HT4 RSMS was correlated with glutamate (R=0.357, p=0.048).

**Conclusions:** We demonstrate glutamate-driven hippocampal remodeling in FEP through a receptor-density gated mechanism, thus providing a mechanistic explanation of how redox dysregulation affects brain structure and symptomatic heterogeneity in schizophrenia.

## Introduction

The hippocampus is a brain structure widely implicated in neurological and psychiatric disorders(1). As part of the memory circuit with several intricate subregions folded onto itself, it remains a key structure that is widely studied for its varied roles in cognition--namely memory, spatial navigation, and perception(2, 3). Beyond the subfields, its functional organization is reputed to follow a “long axis” gradient of specialization with the anterior (or ventral) responsible for anxiety-like behavior, emotion and stress(4–6), and posterior (or dorsal) hippocampus for cognitive functions(6). In conditions such as schizophrenia, the hippocampus is a putative biomarker with early atrophy of select subfields such as the CA1 and subiculum with extension to the rest later in the disease(7, 8).

The hippocampus has been well studied in first-episode psychosis (FEP) and schizophrenia with consistent reports of volumetric reductions across a number of human neuroimaging studies. The consensus from the literature demonstrate an anterior preference and the CA1 and subiculum subfields to be most frequently affected (9–13) while more recent work using higher field MRI show CA4/DG as the earliest to be affected(14). The anterior hippocampus is thought to be the primary site of pathology in schizophrenia given that lesions in the region cause increased dopaminergic transmission and model features of schizophrenia(1, 4, 15), while there is also evidence for abnormalities in the posterior hippocampus(4). In general, studies have focused on the grey matter content of the hippocampus--however, the hippocampus is covered by bundles of white matter that form projections to the rest of the brain. These white matter projections include the alveus, fimbria, and fornix that envelop the hippocampal formation(16), and are relatively understudied in literature with mixed evidence towards their involvement in psychosis(17–19).

Regarding the mechanism for hippocampal degeneration in psychosis, excess glutamate is thought to be pivotal in excitotoxic damage to the brain(7). Elevated levels of glutamate are described in early schizophrenia(20–23) and correlated with reduced grey matter volume(24). This aggregate of glutamate-structural deficit is thought to underlie cognitive and functional impairment(24, 25). Glutamate has been shown to stimulate a hypermetabolic state in the hippocampus preferentially in the CA1 and subiculum during psychosis, followed by early atrophic changes(7). It was hypothesized that the CA1 and subiculum are particularly vulnerable due to the relatively high density NMDA and AMPA receptors compared to other subregions(26), thereby increasing its susceptibility to the excitotoxic effects of glutamate(7). However, this hypothesis is not yet explored in-vivo which we sought to examine in this study.

Here, we aimed to address these gaps in literature by extending the study of hippocampal subfields in FEP to include the white matter subregions, while probing the glutamate-dependent nature of hippocampal atrophy in FEP. First, we sought to replicate previous volumetric studies in FEP examining the hippocampal subfields and shape, while interrogating the entire hippocampal circuit including the white matter tracts. We then tested whether neuroanatomical anomalies would correlate with glutamate receptor densities, by leveraging an in-vivo atlas of the serotonin receptor system(27) combined with gene expression data from the Allen Human Brain Atlas(28) as a method of exploring mutative mechanistic explanations behind structural deficits in FEP. We searched for gene expression correlations between serotonin and glutamate receptor genes such that the distribution of a serotonin receptor subtype may predict glutamate receptor distribution. We tested this hypothesis at the level of the individual by aligning individual hippocampal morphometric and receptor maps, and testing whether the degree of structure-receptor correlation relates to MRS measures of glutamate. We hypothesized that the CA4/DG and CA1 would be amongst the most strongly reduced in FEP, in line with previous studies. Amongst the white matter tracts, the fornix has the strongest evidence for volumetric changes and we expected to replicate this finding(17–19, 29). Shape analysis would likely show contraction in the anterior hippocampus. We hypothesized that hippocampal subfields with higher concentrations of glutamate receptors would show the greatest volumetric differences in FEP.

## Materials and Methods

### Participant and data acquisition

Methods detailing participant recruitment, acquisition of structural and MRS brain imaging are available in Supplementary Methods. Briefly, healthy control (HC) subjects and FEP participants were recruited with lifetime antipsychotic treatment of less than 14 days. All subjects underwent structural brain imaging at 7T resolution, and glutamate levels were estimated using 1H-MRS. The MAGeT Brain algorithm was used for hippocampal morphometry(16, 30–32) to measure subfield volumes and measures of shape(33–35)—details are available in Supplementary Methods.

### Statistical analysis

Multiple linear regression with total hippocampal GM and WM volumes as dependent variables, examining the main effect of diagnosis accounting for age, gender as covariates. Volumetric analyses were conducted at two levels of granularity: using the summed volumes of hippocampal grey matter (GM) and white matter (WM), then at the level of the individual subfields. GM included the CA1, subiculum, CA4DG, CA2CA3, and the SR/SL/SM (stratum radiatum/stratum lacunosum/stratum moleculare; termed stratum henceforth), and WM included the alveus, fimbria, fornix, and mammillary bodies.

### Mapping hippocampal serotonin receptor distributions

We used a previously published atlas of the serotonin receptor subtypes (5-HT1A, 5-HT1B 5-HT2A, 5-HT4) and the serotonin transporter (5-HTT)(27) derived from PET images of 210 healthy subjects to examine 1) Average density of each serotonin receptor within hippocampal subfields, and 2) Density of serotonin receptor subtypes mapped onto hippocampal surfaces, with the aim of correlating structural differences observed between HC and FEP with serotonin receptor densities. Generation of the serotonin atlas is described in the original work--briefly, 210 healthy control subjects underwent PET scanning with 5 tracers to quantify brain-wide density of the serotonin receptors. PET data were collected at approximately 2mm resolution with high-resolution structural imaging at ~1mm for accurate mapping and normalization. The subject maps were aligned to the MNI152 template and averaged to produce a map of serotonin receptor densities at 1mm isotropic voxels. The atlas demonstrated high correlations with postmortem data as well as mRNA expression using the Allen Human Brain Atlas(27).

We generated a MAGeT Brain segmentation of the hippocampal subfields and surface models on the MNI152 template by using the same template library used for segmentation of the subjects (as described above). We inspected the automated segmentation for accuracy prior to measuring average serotonin receptor densities per hippocampal subfield, and further mapped densities to the hippocampal surfaces generated for the MNI152 template. We tested for correlations between 1) Effect sizes quantifying volumetric difference with hippocampal subfields (-log10 of p value) and shape differences (t-statistic), and 2) Serotonin receptor densities at the level of the subfields (average density per subfield) and on hippocampal surfaces.

### Gene expression analysis between serotonin and glutamate receptor genes

The Allen Human Brain Atlas (AHBA) was used to quantify gene expression correlations between serotonin receptor subtypes (5-HT1A, 5-HT1B 5-HT2A, 5-HT4) and 5-HTT, and the glutamate receptor subunits for NMDA and AMPA (Supplementary Table 1). This approach sought to examine which of the serotonin receptors, measured in-vivo using PET, might predict expression of glutamate receptors based on mRNA correlations. Brain gene expression data was obtained from the AHBA database (https://human.brain-map.org/static/download), consisting 6 healthy donor brains. Donors 10021 and 9861 had data for both the left and right hemispheres, while the other 4 donor brains were sampled in the left hemisphere only. Similar to previous work, we chose to only examine the left hemisphere data to ensure consistency in the sampling and to reduce noise(36). We selected only the hippocampal structures sampled in the AHBA which include the CA1, CA2, CA3, CA4, DG, subiculum, mammillary body, as well as the medial and lateral mammillary nucleus separately.

For genes with multiple probes, the optimal probe was chosen based on quality control data previously published(37). This was based on comparison of gene expression data between the AHBA Agilent microarray and the “ground truth”data from RNA-Seq, where the correlation between each probe and matching gene in RNA-Seq was measured and the probe with the greatest correlation to RNA-Seq was determined as the best probe. A probe for a gene was chosen if it passed quality control (is_passing_probe=TRUE, probes with q < 0.1 based on correlation with RNA-seq data), and filtered based on the highest correlation to RNA-seq data, as denoted in the “is_best_probe” column (highest correlation). We opted for this procedure of selecting only one probe per gene, as genes had varying number of probes and taking the average may imbalance gene weights in downstream analysis. This step yielded one probe sampling one gene per 6 left hemispheres. The median expression per gene was then calculated across 6 brains, followed by normalization across structures so that the values are comparable across genes.

Our genes of interest included all of the known subunits for NMDA and AMPA receptors, as well as the serotonin receptor subtypes--adding up to 15 in total (Supplementary Table 1). All genes were available in the quality controlled AHBA data except GRIN3B as it is a more recently discovered subtype. Gene-gene Pearson’s R correlations were visualized, and we examined whether certain serotonin receptor genes clustered together with the glutamate receptor genes.

### Reproducibility using an alternate definition of hippocampal subfields

To test the reliability of our results using a different automated segmentation algorithm, the T1-weighted MRIs were additionally processed using the FreeSurfer hippocampal subfield module (http://surfer.nmr.mgh.harvard.edu/fswiki/HippocampalSubfields)(38). The procedure stems from ultra-high resolution, ex-vivo MRI based definitions of the hippocampal subregions at 0.13mm isotropic resolution acquired from fifteen autopsy samples. A computational atlas was then compiled from the manual labels, released as part of a module within the FreeSurfer software package (version 6.0), and well validated for use with in-vivo MRI. Our subjects as well as the MNI152 template were processed through the FreeSurfer software to generate an independent set of hippocampal subfield volumes and average serotonin receptor densities. Standard settings were applied, with resulting outputs visually inspected using the same procedure for MAGeT Brain hippocampal segmentations.

### Reproducibility across different samples

We further tested the reliability of subfield-to-receptor correlations using data from published studies examining hippocampal subfields in psychosis. To this end, we selected three recent studies with Studies 1 (41 FEP, 39 HC(39)) and 2 (58 FEP, 76 HC(40)) examining first-episode psychosis, and the third study with chronic schizophrenia (155 SCZ, 79 HC(8)) and mean duration of illness was 7 years. Samples in Studies 1 and 2 had minimal antipsychotic exposure--Study 1 was medication naive, while Study 2 excluded patients with antipsychotic treatment greater than 3 months. Study 3 was chosen to compare findings in a FEP population vs. chronic schizophrenia, with the hypothesis that structure-receptor correlations in chronic schizophrenia would likely reflect medication changes while those in FEP would more closely follow their underlying neurobiological changes linked to glutamate. The mean daily dose was 212.3 (CPZ equivalent) for Study 3.

All three studies used the same hippocampal subfield module within FreeSurfer. Studies 1 and 2 used version 6.0 while study 3 used version 5.3, while both versions utilize the same computational atlas with minor differences in outputs(38). We extracted p-values of subfield volumetric comparisons between healthy controls and the psychosis groups, which were -log10 transformed. Studies 2 and 3(8, 40) reported findings from select subfields including GC-ML-DG, CA1, CA3, CA4, molecular layer, hippocampal tail, and presubiculum (only for Study 2) while Study 1 report on all subfields. Therefore, we restricted further analyses for all three studies within the 8 subfields stated above. We repeated the correlations between subfield effect sizes and serotonin receptor densities. We expected to replicate findings, specifically with strongest correlations for 5-HT1A and 5-HT4, Studies 1 and 2 but not Study 3.

### Normative modelling to examine individual morphometric patterns

To examine whether the structure-receptor correlations translate to the level of the individual, we generated normative maps of hippocampal shape. Hippocampal shape measures were first regressed against age and sex in the entire cohort (including HC and FEP). The residuals for the FEP group were then z-scored against the HC mean and standard deviation at each vertex. The resulting z-score map is then correlated against the distribution of 5 serotonin receptors on the hippocampal surface, yielding 10 correlations (5 per hemisphere) which we term Receptor-Specific Morphometric Signatures (RSMS). We explored whether RSMS was correlated with individual measures of ACC glutamate, restricting our analysis to 5-HT1A and 5-HT4-RSMS. We hypothesized that RSMS measures would positively correlate with glutamate, further supporting glutamate-dependent model of hippocampal remodelling that follows the distribution of glutamate receptors.

## Results

The final sample consisted of 68 subjects (27 HC, 41 FEP) passing image preprocessing quality control. 18 subjects (20.9% of initial cohort) were excluded due to errors in automated hippocampal analysis. Group-wise demographic data are available in Supplementary Table 2--there were no significant differences in distribution of age or sex distribution between the groups.

There were volumetric differences observed at the level of the whole GM and WM between HC and FEP (Figure 1a). Both left (t= −2.22, p=0.030, DF= 64). and right (t= −2.552, p= 0.013) hippocampal GM were significantly smaller in FEP (Figure 1a). The WM was initially did not differ significantly on the left (t=-1.51, p= 0.137), but did so for the right (t=-2.289, p= 0.025) (Figure 1b). After excluding one outlying HC subject for left WM (based on volume less than 3 SD below the mean & visual inspection of data), both left (t=-2.425, p=0.0182, DF=63) and right (t= −2.402, p=0.0193) WM were reduced in FEP. At this level of anatomical granularity--multiple testing correction (Bonferroni) for 4 comparisons (two hemispheres x WM and GM) gives P value threshold of p < 0.0125 for significance after correction, meaning none of the findings would survive after correction. At the level of the subfields, the CA4-DG show greatest difference amongst subfields, with both the left (t=-2.70, p=8.91E-03) right (t=-2.87, p=5.50E-03), reduced in FEP compared to HCs (Figure 1c).

Vertex-wise analysis of shape differences between HC and FEP did not yield any findings that survived FDR correction. However, examining subthreshold vertices shows contraction of the bilateral anterior hippocampus (Figure 1d). T-statistics ranged from −2.715 to 2.154 for the left, and −3.103 to 1.906 for the right. At the peak vertex in the right anterior hippocampus, there is significant contraction in FEP compared to HC (t= −3.103, p= 2.85E-03, DF=64) (Figure 1d).

**Figure 1.**
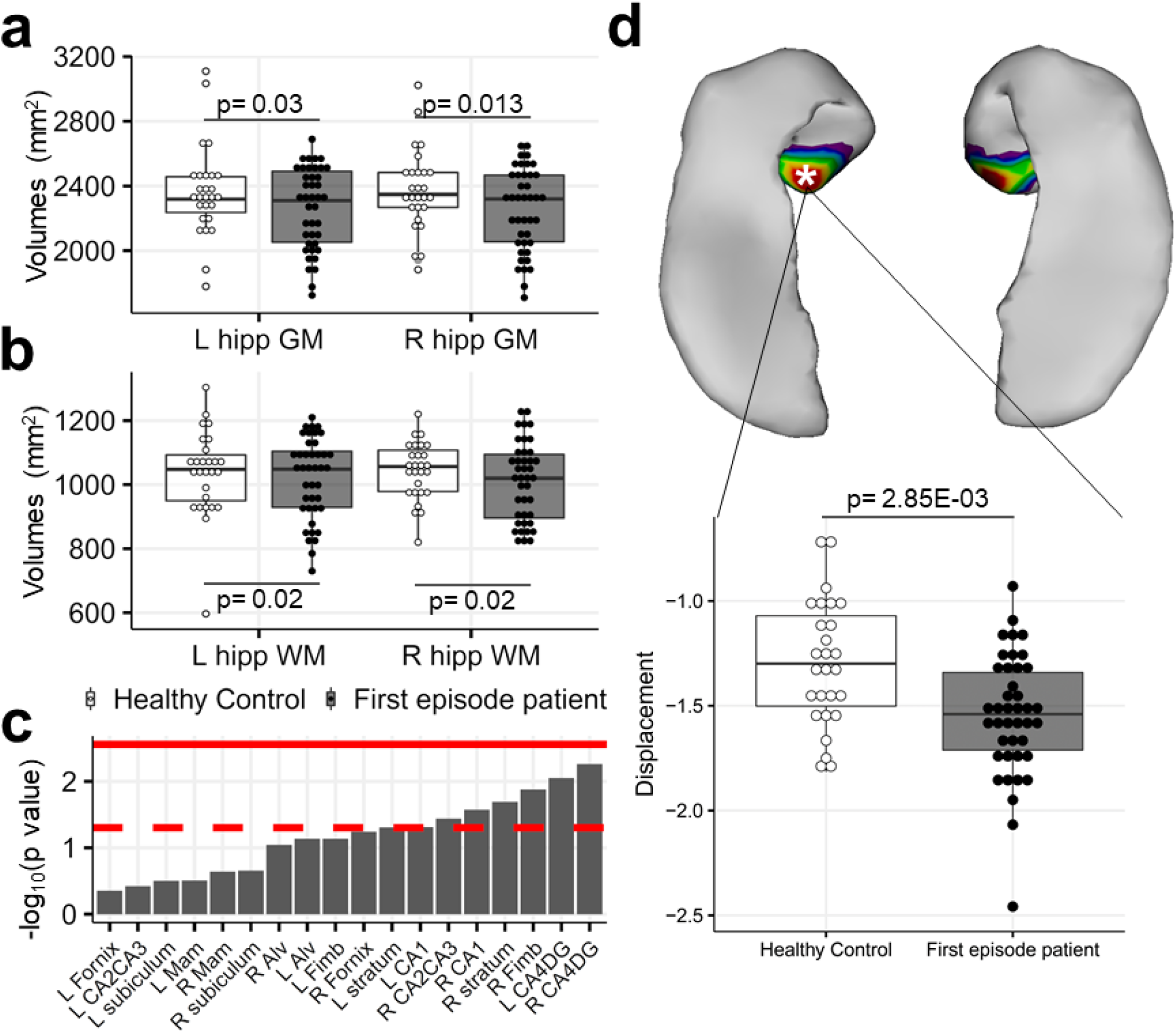
Volumetric and shape analysis of the hippocampus in FEP compared to healthy controls. a-b) Distribution and comparison of hippocampal GM and WM volumes. P-values for left hippocampal WM are after excluding 1 outlying HC subject. c) Subfield-wise comparisons between HC and FEP. Solid red line indicates the Bonferroni-corrected p-value threshold for 18 subregions, while the dotted red line indicates p=0.05. d) Shape analysis demonstrating bilateral contraction of the anterior hippocampus in FEP compared to HC.

Next, we examined gene expression correlations to look for serotonin receptors that might best predict glutamate receptor expression (Figure 2a). Based on the gene-gene correlations in the AHBA, we found the serotonin receptor genes HTR1A (encoding the 5-HT1A subtype) and HTR4 (encoding the 5-HT4 subtype) to be closely associated with glutamate receptor genes based on Pearson’s correlations (Figure 2b). HTR1A was highly positively correlated with GRIA1-3 (AMPA subunits), and HTR4 with GRIN1, GRIN2A-C (NMDA subunits) (Figure 2b). Therefore, we predicted the 5-HT1A and 5-HT4 receptor distributions, which reflect AMPA and NMDA glutamate receptor densities, to demonstrate strongest correlations to volumetric differences. In addition, we found SLC6A4 (5-HTT) to negatively correlate (with Pearson’s R ~0.50) with most glutamate receptor genes (Supplementary Figure 1). We estimated serotonin receptor densities on the hippocampal surface and subfields by generating MAGeT Brain segmentations on the MNI152 template (Figure 2c) (Supplementary Figure 2).

**Figure 2.**
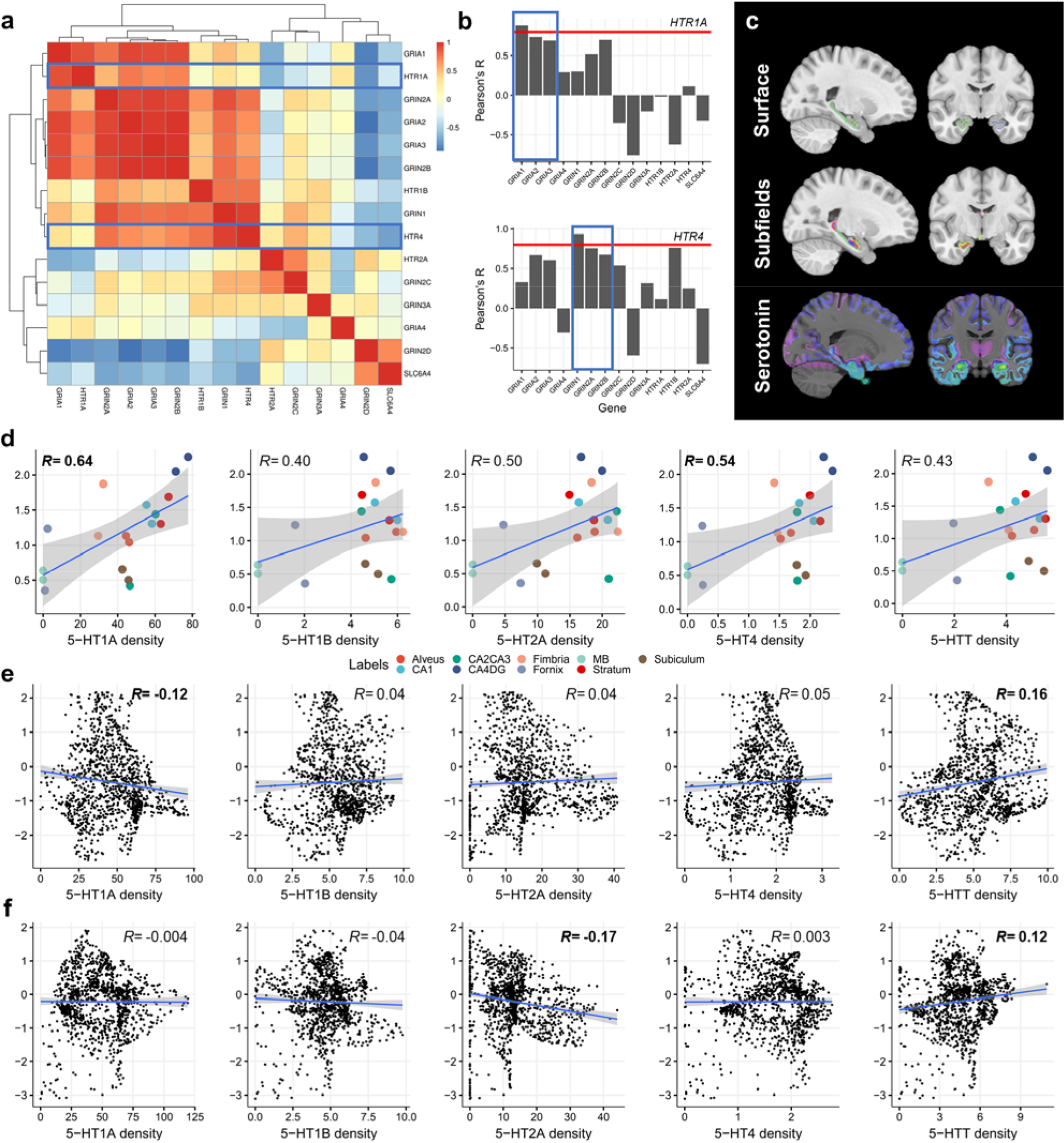
Linking serotonin and glutamate receptors through gene expression analysis, and then to volumetric differences. a) Gene expression correlations between 13 genes sampled in the AHBA for AMPA, NMDA subunits and serotonin receptor subtypes. Blue boxes are 2 candidate genes (HTR1A and HTR4) for predicting glutamate receptor expression. b) HTR1A and HTR4 expression was significantly correlated with GRIA1-3 and GRIN1-2 genes (outlined in blue) that code for AMPA and NMDA receptor subunits respectively. Red line indicates Pearson’s R= 0.80. c) MNI152 template with the hippocampal surface and subfields outlined using MAGeT Brain, which are then combined with the serotonin receptor atlas to yield receptor densities across each hippocampal surface and average density per subfield. d) Correlations between effect sizes (HC vs. FEP) measured as −log10(p value) and average receptor densities across the hippocampal subfields. e-f) Correlations between hippocampal shape effect sizes (HC vs. FEP) measured as t-statistics and receptor densities per vertex, for the left (e) and right (f) hippocampus.

We tested correlations between 1) Subfield effect sizes resulting from multiple linear regression (-log10 of the p value), and 2) Mean density of serotonin receptor subtypes. As predicted, 5-HT1A showed the strongest correlation (Pearson’s R= 0.64, p= 4.09E-03, and the second strongest correlation was 5-HT4 (R= 0.54, p= 0.02). 5-HT1B, 2A, and 5-HTT showed modest positive correlations (R= 0.40, 0.50, 0.43, and p= 0.10, 0.03, 0.07 respectively (Figure 2d). We repeated the structure-receptor density correlation analysis using t-statistics from hippocampal shape analysis, which revealed similar findings. For the left hippocampus, 5-HT1A density demonstrated the strongest correlations to structural effect sizes (R= −0.119, p= 4.827E-05), followed by 5-HTT (R= 0.161, p=3.658E-08) (Figure 2e). 5-HT4 was not significantly correlated (R= 0.050, p= 0.0913), as was 5-HT1B and 2A (R= 0.039, 0.042, p= 0.182, 0.154) (Figure 2e). The right hippocampus showed somewhat contrasting findings of 5-HT2A density correlating significantly (R= −0.168, p= 3.723E-09) as well as 5-HTT (R= 0.120, p= 2.564E-05), with 5-HT1A, 1B, and 4 not well correlated (R= −0.004, −0.038, 0.003, p= 0.886, 0.185, 0.920) (Figure 2e).

We repeated our volumetric analysis and subfield effect size-receptor correlations using a different hippocampal subfield analysis pipeline through FreeSurfer. 61 subjects (38 FEP, 23 HC) passed quality control (Supplementary Figure 3). Volumetric analysis demonstrate the left (t=-2.05, p=0.045) and right (t=-2.74, p=8.26E-03) hippocampi to be reduced in FEP (Supplementary Figure 4). FreeSurfer subfields analyses show the R CA1 (t=-2.90, p=5.24E-03), molecular layer, hippocampal tail, and fimbria with the strongest effect sizes (Supplementary Figure 4). None of the subfield results survive multiple testing correction. In repeating the subfield-serotonin correlations, we partly reproduce our results with 5-HT1A (R= 0.372, p= 0.074) and 5-HT4 (R= 0.292, p= 0.167) demonstrating strongest correlations to effect sizes, while the other receptor subtypes had weaker correlations in comparison (Figure 3, top row, and Supplementary Table 3). We then sought replication across different samples examining hippocampal subfields in psychosis using data from 3 recently published studies. Here, we demonstrate replication of our hypothesis with the hippocampal subfield effect sizes correlating most strongly with the distribution of 5-HT1A and 5-HT4 as in our own sample--this was evident for Study 1 (5-HT1A: R=0.78, p= 3.65E-04, 5-HT4: R= 0.66, p= 5.80E-03) and Study 2 (5-HT1A: R=0.68, p= 3.82E-03, 5-HT4: R= 0.60, p= 0.0138) (Figure 3, blue and green boxes). Study 3 effect sizes showed significant correlations for 5-HT1B (R= −0.57, p= 0.035) and 5-HT2A (R= −0.65, p= 0.011), but not for the other receptor subtypes as expected. Full table of Pearson’s correlations and p-values are available in Supplementary Table 3.

**Figure 3.**
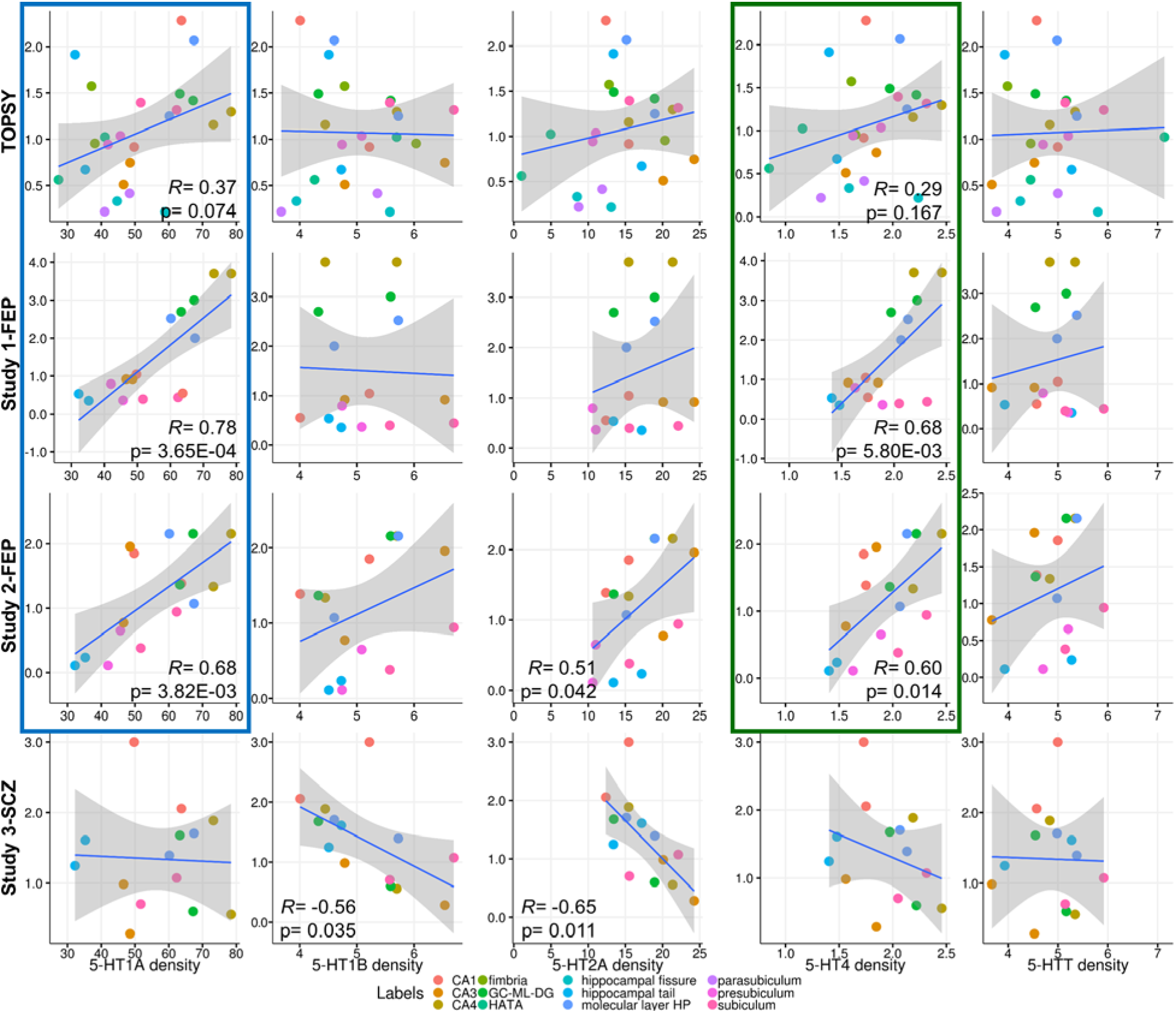
Correlations between effect sizes measured as −log10(p value) on the y-axis and average receptor densities across the hippocampal subfields as defined by FreeSurfer. Studies 1 and 2 consisted of data from FEP subjects while Study 3 studied chronic schizophrenia. Effect sizes from our cohort and Studies 1-2 all demonstrate strongest correlations with 5-HT1A density (blue box) and 5-HT4 (green box), consistent with the hypothesis.

Lastly, we used normative maps of hippocampal shape to generate individuals measures of structure-receptor alignment, termed RSMS (Figure 4a). Amongst the FEP group, 31 patients had MRS data available. Testing for correlations between 5-HT1A and 5HT4-RSMS with MRS measures of glutamate showed significant correlation between the right 5-HT4-RSMS with glutamate (R= 0.357, p= 0.048) (Figure 4b). The left 5-HT4-RSMS was not significantly correlated with MRS measures, nor was 5-HT1A-RSMS (p > 0.10).

**Figure 4.**
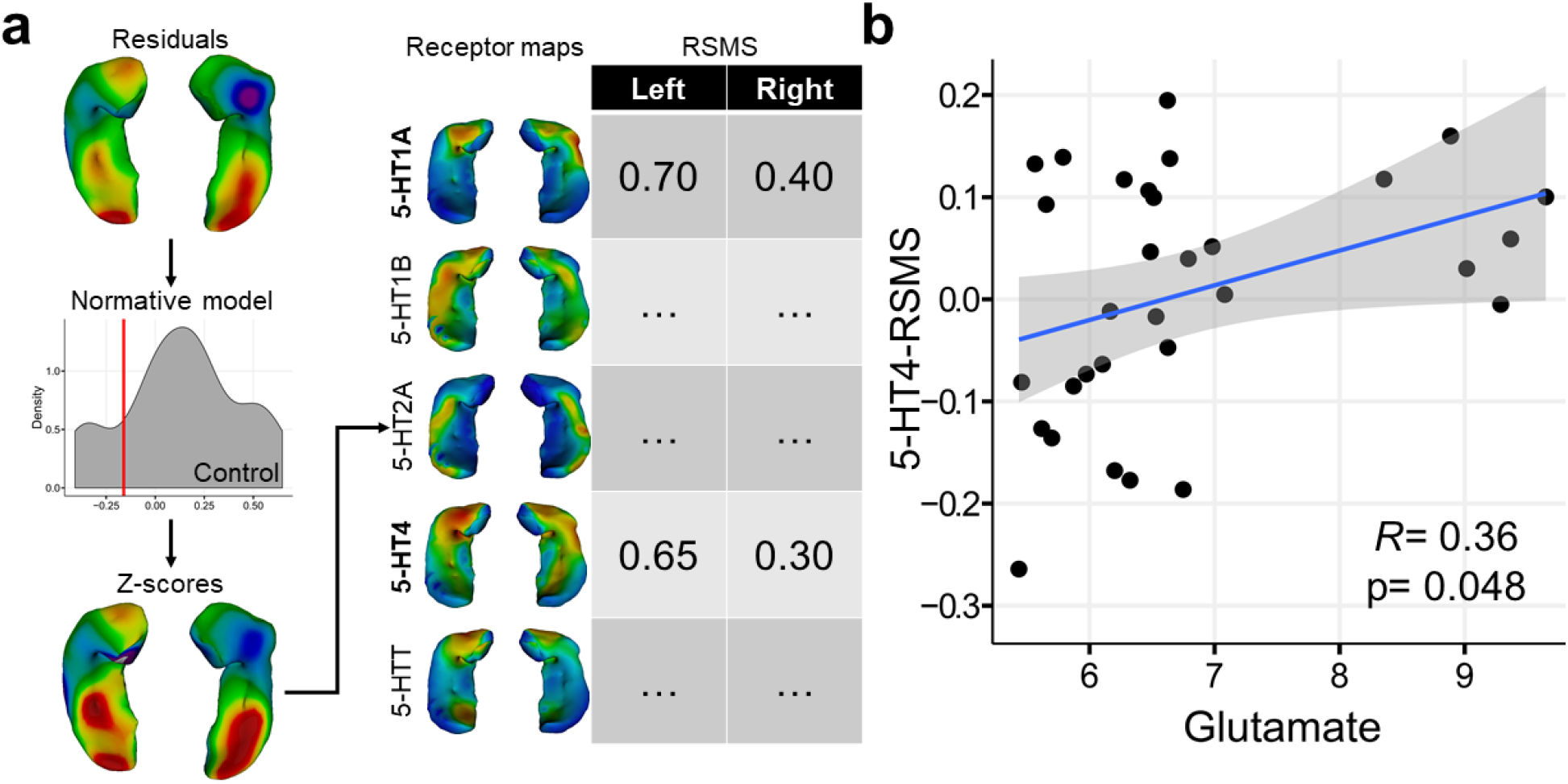
Normative modelling of hippocampal shape to derive Receptor-specific Morphometric Signatures (RSMS) and correlations to MRS measures. a) Illustration of normative modelling and RSMS generation. Shape measures are first regressed against age and sex to yield residuals for the entire cohort. Each FEP subject is then z-scored against the HC distribution per vertex, resulting in a normative map of shape measures. The z-scores are correlated with each of the serotonin receptor maps per hemisphere, resulting in 5 RSMS values per hemisphere. b) Significant positive correlation between MRS measures of glutamate with right-sided 5-HT4-RSMS.

## Discussion

Neuroimaging studies in schizophrenia have long since demonstrated structural changes in early schizophrenia. Here, we replicated previous findings of regional selectivity in volume loss within FEP patients. For the first time, we linked hippocampal structural changes in psychosis to the distribution of serotonin receptors 5-HT1A (at the subfield and vertex level) and 5-HT4 (at the subfield level), thus by proxy AMPA and NMDA receptors. We replicated this association using published data from two FEP studies with respect to 5-HT1A and 5-HT4 specific correlations at the level of subfields. Lastly, we demonstrate that higher MRS levels of glutamate is associated with 5-HT4-specific morphometric signatures of hippocampal shape, and thus suggestive of NMDA-mediated morphometric changes. Moreover, by capturing the concordance between brain structural profiles and receptor densities, we demonstrate that excitotoxicity-related structural damage becomes less prominent in chronic stages of schizophrenia. Apart from providing first hand proof for the glutamatergic theory of morphological changes in schizophrenia, these results uncover receptor distribution as a critical mediator of the interindividual heterogeneity in the neuroanatomy of schizophrenia.

In our structural analyses, we replicate previous findings of reduced hippocampal volumes bilaterally, without significant asymmetry in findings (Figure 1). Interestingly, the WM also demonstrated similar reductions bilaterally. At the level of subfields, we found the CA4/DG subfield to be the most affected bilaterally in FEP--in line with more recent studies using high-resolution imaging(14), while other studies using standard resolution generally report CA1 as the key subfield. Methodological differences likely contribute to this discrepancy between findings with the choice of hippocampal subfield delineation affecting outcomes. For example, our own FreeSurfer-based analysis showed CA1 to be the most affected in FEP rather than CA4/DG. The definitions of the hippocampal subfields differ between the two automated methods (MAGeT Brain and FreeSurfer) used in this study as well as the computational aspects of the algorithms themselves, highlighting the caution needed in interpreting the sequential structural changes in the subregions (for instance, the differences among Lieberman(41) (primacy of CA1); Tamminga(42) (primacy of CA3) and Harrison(43) (primacy of CA4)). However, there is a notable consistency across methods (Freesurfer, MAGet parcellations, and vertexwise surface analysis) in localizing the FES-specific changes to the anterior hippocampus and subregions more prominent in the anterior hippocampus such as the CA1, CA4/DG. The CA1 and CA4/DG are tightly folded together in the anterior hippocampus and it is possible that changes in either subfield could be attributed to any one(31). Another significance of our findings is that we were amongst a few studies that studied medication-naive FEP patients. A brief literature search in FEP and hippocampus shows that only 2 out of 22 studies (Supplementary Table 4) could be considered as having medication-naive FEP patients. Given antipsychotic medications alter brain structure, our cohort with 1.45 median DDD days of antipsychotic exposure contributes a clearer perspective towards grey matter changes in early psychosis. We also highlight the hippocampal white matter that showed reductions as a whole (Figure 1b) and within subregions including the fimbria, alveus and fornix (Figure 1c), in keeping with the histopathological studies of Heckers et al(44) and Eastwood et al(45) as well as the neuroimaging observations(17, 18, 29, 46). These were not unexpected findings given that the grey and white matter subfields form closely related pathways originating from the entorhinal cortex(16), and are often impacted in a prion-like progressive pathology along the circuitry(1, 16, 47). This prompts a reconsideration of the mechanistic basis of the putative progressive volumetric changes in hippocampus in schizophrenia.

We found 5-HT1A to correlate with AMPA subunit expression and 5-HT4 with NMDA subunits based on gene expression analysis. Both 5HT1A and 5HT4 are preferentially expressed during early embryonic development(48), prompting a developmental interpretation of the hippocampal volume deficits in schizophrenia. Further, both 5-HT1A(49) and 5-HT4(50, 51) expression is enriched in the ventral pole (anterior hippocampus)(51), and the dentate gyrus(52). In line with this, higher NMDA and AMPA expression has been reported in the anterior CA1 than CA3(26, 53). There is evidence for co-regulation of 5-HT1A, AMPA, as well as NMDA receptor expression in the hippocampus(54) which support our findings. Further, 5-HT1A and AMPA are tightly coupled in that 5-HT1A antagonist increased AMPA subunit levels(55), while 5-HT1A modulates AMPA currents(56), indicating a high likelihood for neuroanatomical co-localization. 5-HT4 receptor functions to increase the excitability of hippocampal pyramidal neurons, by reducing long-term depression (LTD), a key NMDA-mediated synaptic plasticity mechanism(57). 5HT4 receptors play a crucial role in the potentiation of pyramidal CA3-CA1 synapses of the trisynaptic pathway(58), and promote dendritic spine formation(52), explaining the observed coexpression with NMDA receptors. Taken together, our results are in line with the strong evidence for serotonin and glutamate receptor co-localization within the hippocampus, explaining the increased susceptibility to glutamate-mediated changes in CA1 and dentate gyrus, and the ventral hippocampal surface.

We confirmed the pattern of morphometric abnormalities in the hippocampus correlate with the distribution of glutamate receptors through serotonin receptor proxies, consistent with our initial hypothesis. By using the 5-HT1A receptor distribution as a proxy for AMPA and 5-HT4 for NMDA receptor expression, we show positive correlations between receptor density and subfield effect sizes--meaning that regions with greatest amount of volume loss in FEP have higher glutamate receptor expression. We partly replicate this finding within the study sample using an alternative definition of the hippocampal subfields using the FreeSurfer software, with 5-HT1A and 5-HT4 as the most significant correlations to effect sizes (Figure 3). Our hypothesis is further supported by consistent replication based on data from two additional FEP studies(39, 40), and that these findings are specific to early psychosis rather than chronic schizophrenia (Figure 3). Within our own sample, we did not fully replicate the structure-receptor correlations using FreeSurfer--this may be due to the FreeSurfer definition of hippocampal subfields being much smaller than the Winterburn atlas(31) used in MAGeT Brain, thereby impacting the accuracy of serotonin receptor sampling at a resolution of 1mm. Vertex-wise correlations between shape differences and receptor densities were somewhat less consistent than our subfield findings likely due to the receptor distributions only applying to the outer surface of the hippocampus. For example, volumetric changes occurring for subfields embedded within the hippocampus would not necessarily reflect on the surface-level findings which may contribute to some level of inconsistency in structure-receptor correlations on the vertex-wise level. Although surface-based sampling allows for finer sampling of receptor densities across a continuous surface, structural changes that occur below the surface may not be captured accurately which poses a limitation.

In testing the subfield-receptor correlations using published results from a chronic schizophrenia population, we found that structural anomalies correlated with 5-HT1B and 5-HT2A with a reversal of correlations--subfields with higher expression of receptors had less pronounced structural deficits. Contrasting these with findings in FEP populations, this hints at structural changes secondary to antipsychotic medications particularly considering that neuroleptics tend to antagonize 5-HT1B and 5-HT2A(59, 60). We speculate possible neuroprotective effects of 5-HT2A antagonism as chronic 5-HT2A blockade was shown to promote hippocampal neurogenesis(61). Further, antipsychotics reduce NMDA receptor availability in CA1 and CA2 the long term(62), which may contribute to the correlation reversal. Although not the focus of the current study, this remains an important future avenue for investigation.

Extant neuropathological evidence indicates that a reduction in dendritic spines as well as the number of GABAergic interneurons, rather than a reduction in pyramidal cell numbers, is the likely substrate of hippocampal volume reduction in schizophrenia(63). In this context, two competing interpretations could be made from our observations. Functionally, interneuron reduction may result in a disinhibition effect, with consequent high glutamatergic output at mPFC synapses, explaining the observed relationship between ACC glutamate levels and hippocampal volume changes in regions expected to have a higher NMDA expression. This notion is supported by ketamine related effects in CA1 and further propagation of hypermetabolism with repeated glutamatergic surge observed in mice(7). Alternatively, reduced dendritic spines in anterior hippocampus may be secondary to prefrontal glutamatergic excess leading to excitotoxic damage, and may serve to compensate for the resulting hypermetabolism in hippocampus.

We sought to further support these group-level findings using individual measures. Through normative modelling and assessing individual morphometric patterns and their alignment to the serotonin receptor maps, we further illustrate a glutamate-dependent mechanism for hippocampal atrophy. The individual measures of alignment likely provide a more accurate measure than group differences at the level of subfields since there is greater sampling (~1100 vertices per hemisphere) for accurate receptor density measures, and thus the correlations are more reliable. Right 5-HT4 RSMS, indicative of the resemblance of an individual’s structural profile to that of the 5-HT4 (NMDA) receptor distribution, was positively correlated with MRS measures of glutamate (Figure 4b). This suggests that structural changes in the right hippocampus, specifically, is more closely related to excess glutamate through an NMDA receptor-dependent mechanism. Given the lack of relationships between left-sided RSMS and glutamate, changes in the left hippocampus may be due to a separate mechanism or due to limitations with surface-based modelling. A limitation may be that MRS measures are measured the dorsal ACC and not in hippocampus, and the correlations may have been stronger if glutamate values from the hippocampus were available. Nevertheless, given the strong monosynaptic as well as indirect projections between the medial prefrontal cortex and ACC with the hippocampus(64), the ACC MRS glutamate concentration may indeed reflect the hippocampal-prefrontal synaptic excitatory tone, explaining the observed relationship with hippocampal shape. Beyond glutamate, other systems including dopamine and serotonin may be involved, as neurotransmitters such as glutamate, dopamine, and serotonin have complex 3-way relationships—for example, glutamate dysfunction is linked to dopamine synthesis and psychosis(4, 15, 65), serotonin metabolites are increased in FEP(66), while clarifying this fully is beyond the scope of this work(67).

We note several limitations of this study, including the relatively small sample size and the cross-sectional analyses. The size of both our study groups (FEP and HC) were limited due to accessing medication-naive FEP individuals (max treatment 3 days of antipsychotics), with matched healthy controls based on parental education. While our sample was closely matched with respect to age, sex, and socioeconomic status, this restricted our power to detect significant differences. The automated methods for analyzing the hippocampus and subfields is a potential source of discrepancy--however we attempt to show consistency between two different algorithms (MAGeT Brain and FreeSurfer) in comparing our results to previous studies.

By highlighting the role of 5HT1A and 5HT4 in glutamate-mediated hippocampal volume reduction in psychosis, our study raises the question of repurposing or employing existing agents with preferential 5HT1A (e.g. lurasidone, cariprazine) and 5HT4 affinity (e.g. prucalopride) as neuroprotective agents in schizophrenia. Hippocampal morphometry may provide the much-needed treatment engagement targets for the putative procognitive agents affecting 5HT1A and 5HT4 system. We advocate for the use of structural MRI, as well as MRS in clinical trials evaluating such agents, to generate the conclusive experimental evidence for the glutamatergic hypothesis of hippocampal dysfunction in schizophrenia.

## Acknowledgements

Computations were performed on the Niagara supercomputer at the SciNet HPC Consortium (Loken et al., 2010). SciNet is funded by: the Canada Foundation for Innovation; the Government of Ontario; Ontario Research Fund - Research Excellence; and the University of Toronto.

We thank Dr. Joe Gati, for their assistance in data acquisition and archiving. We thank Dr. William Pavlovsky for consultations on clinical radiological queries. We thank Drs. Raj Harricharan, Julie Richard, Priya Subramaniam and Hooman Ganjavi and all staff members of the PEPP London team for their assistance in patient recruitment and supporting clinical care. We gratefully acknowledge the participants and their family members for their contributions. The MRS pulse sequence package used was developed from a source code provided by Uzay E. Emir, PhD, and Dinesh K. Deelchand, PhD, Center for Magnetic Resonance Research (CMRR), Minneapolis, MN. It is based on original work by Gülin Öz and Ivan Tkáč(68). Our special thanks to Uzay E. Emir, PhD, Dinesh K. Deelchand, PhD, Gülin Öz, PhD, Ivan Tkáč, PhD, and Edward Auerbach, PhD, for their help with sequence testing and for valuable technical discussions and to Dennis W. J. Klomp, PhD, and Vincent O. Boer, PhD, for providing the asymmetric excitation pulse used in the 7T version of the sequence. We also acknowledge CFREF BrainsCAN funding that we received providing a reduced hourly rate for the MRI scans.

